# Quantitative systems pharmacology modeling of avadomide-induced neutropenia enables virtual clinical dose and schedule finding studies

**DOI:** 10.1101/2021.04.28.438168

**Authors:** Roberto A. Abbiati, Michael Pourdehnad, Soraya Carrancio, Daniel W. Pierce, Shailaja Kasibhatla, Mark McConnell, Matthew W. B. Trotter, Remco Loos, Cristina C. Santini, Alexander V. Ratushny

## Abstract

Avadomide is a cereblon E3 ligase modulator and a potent antitumor and immunomodulatory agent. Avadomide trials are challenged by neutropenia as a major adverse event and a dose-limiting toxicity. Intermittent dosing schedules supported by preclinical data provide a strategy to reduce frequency and severity of neutropenia, however the identification of optimal dosing schedules remains a clinical challenge.

Quantitative Systems Pharmacology (QSP) modeling offers opportunities for virtual screening of efficacy and toxicity levels produced by alternative dose and schedule regimens, thereby supporting decision-making in translational drug development.

We formulated a QSP model to capture the mechanism of avadomide-induced neutropenia, which involves cereblon-mediated degradation of transcription factor Ikaros, resulting in a maturation block of the neutrophil lineage.

The neutropenia model was integrated with avadomide-specific pharmacokinetic and pharmacodynamic models to capture dose-dependent effects. Additionally, we generated a disease-specific virtual patient population to represent the variability in patient characteristics and response to treatment observed for a diffuse large B-cell lymphoma trial cohort.

Model utility was demonstrated by simulating avadomide effect in the virtual population for various dosing schedules and determining the incidence of high-grade neutropenia, its duration, and the probability of recovery to low grade-neutropenia.

## Introduction

Neutrophils are a major class of white blood cells (1). Neutrophils mature in the bone marrow, move to and reside in peripheral blood circulation, and migrate to inflamed tissue sites when necessary (2). Here, neutrophils can degranulate, phagocyte microbes, or release cytokines to amplify inflammatory response (3). The blood count of neutrophils (absolute neutrophil count or ANC) is a clinical metric for individual capability to fight infections. Neutropenia is a state of low ANC (4,5), which can occur due to genetic disorders (e.g., cyclic neutropenia), immune diseases (e.g., Crohn’s disease), or may occur as a drug-induced toxicity (6).

IMiDs and CELMoDs are a class of compounds therapeutically active against a number of malignancies. These therapeutics include thalidomide, lenalidomide, pomalidomide (7) and others currently in clinical development (e.g., Iberdomide (8)). IMiD/CELMoD compounds bind to cereblon (CRBN) and modulate the affinity of the cereblon E3 ubiquitin ligase complex (CRL4^CRBN^) to its substrates, thereby favoring their recruitment, ubiquitination and subsequent proteasomal degradation. Avadomide (CC-122) is a novel CELMoD being developed for patients with advanced solid tumors, non-Hodgkin lymphoma (NHL), and multiple myeloma (MM) (9). While research continues towards full elucidation of avadomide activity, it is known that avadomide drives CRL4^CRBN^ interaction with two hematopoietic zinc finger transcription factors Ikaros (IKZF1) and Aiolos (IKZF3) inducing their degradation. These transcription factors are known to promote immune cell maturation (10) and normal B- and T-cell function (11). Avadomide administration is associated with a potent antitumor effect and stimulation of T and NK cells in diffuse large B-cell lymphoma (DLBCL) patients (12).

In a recent phase I trial for avadomide in patients with advanced solid tumors, NHL, or MM (Trial Identifier: NCT01421524), 85% of patients experienced treatment-emergent Grade 3/4 adverse events, primarily neutropenia, followed by infections, anemia, and febrile neutropenia (13). Clinical management of neutropenia includes adjunct therapies to stimulate neutrophil production (e.g., administration of granulocytic-colony stimulating growth factor (G-CSF) as filgrastim), dose-reduction, or treatment discontinuation. Another approach to manage avadomide-induced neutropenia is the introduction of an intermittent dosing schedule. For example, 5 days on-followed by 2 days off-treatment (5/7 schedule) improved tolerability and reduced frequency and severity of neutropenia, febrile neutropenia, and infections (13).

In this context, quantitative systems pharmacology (QSP) modeling offers opportunities for *in silico* exploration of alternative dose and schedules that maximize drug exposure while allowing for toxicity management. Such a QSP tool is much needed because CELMoDs are a large and growing family of compounds and many CELMoDs developed to date share similar patterns of toxicity.

Several authors have published mathematical models of neutrophil maturation and neutropenia state, readers are encouraged to read the review by Craig (14). Some shared characteristics emerge among differential equation based models: (i) the presence of a proliferative neutrophil progenitor pool (15), (ii) sequential maturation stages in bone marrow followed by egress into peripheral blood, (iii) fixed life span of neutrophils in circulation, and (iv) some form of control mechanism that regulates neutrophil level (16–18). Further papers highlight the existence of a reservoir pool of mature neutrophils in bone marrow (19,20) and of a marginated pool of neutrophils (consisting of neutrophils localized in sites other than bone marrow and peripheral blood that are able to relocate) (21,22).

Here, we develop a QSP model to represent avadomide-induced neutropenia and we apply it to predict the incidence and the severity of neutropenic events in a virtual DLBCL (diffuse large B-cell lymphoma) population across a range of dosing schedules to demonstrate its potential utility.

The model development followed relevant good practice guidelines (23,24) and included verification of model structural identifiability (25–27), global sensitivity analysis (28) and model validation (29).

## Methods

This section details technical and methodological aspects of model implementation.

### ODE based models

The models for avadomide-pharmacokinetics (PK) and neutrophil life cycle are ordinary differential equation (ODE) based and were integrated using Matlab R2020a ODE routines (30). For model fit we applied the optimization routine *fminsearch* (31) to minimize an objective function consisting in the weighted sum of absolute normalized difference between model simulation and experimental data.

### Model structural identifiability and global sensitivity analysis

Structural identifiability verifies that, given the proposed model structure, it is possible to regress a unique set of model parameters (globally or locally) under the hypothesis of ideal data (noise-free and continuously sampled) (32). This test was conducted in Matlab using the GenSSI 2.0 package (33–35).

Sensitivity analysis (SA) allows exploration of model input-output structure and supports model development. Global SA (GSA) enables a broad exploration of parameter space. We adopted a Monte Carlo based method as described in (36) (Supplementary Material 1.1).

### Virtual patient population

To represent the heterogeneity of ANC data observed in the clinical trial, we generated virtual patients representing clinical disease-specific cohorts. A virtual patient consists of a neutrophil life cycle model for which selected parameters are assigned from probability functions determining the expected parameter distributions for patients having a given tumor type (e.g., Glioblastoma (GBM) or DLBCL). These probability distribution functions are generated by repeated model fit to individual clinical ANC data, thereby estimating the parameter value empirical distributions. These distributions are tested for normality by applying the Anderson-Darling test (*adtest*, Matlab) and smoothed adopting a kernel density estimation (*ksdensity*, Matlab).

### Model validation

For validation, the model simulations were compared to clinical datasets that were not used during the virtual population development. The comparison was based on a two-sample Kolmogorov-Smirnov (K-S) test. This statistical test determines if the empirical distributions of two sample sets belong to the same distribution. Here, the two sample sets are the model generated ANC and clinical ANC taken at the same time after avadomide administration. This test was executed in Matlab using the *kstest2* function.

### Estimation of toxicity

The final goal of the simulation is the quantification of neutropenia incidence for a given avadomide dosing schedule in a virtual patient population. We focused on neutropenia and did not develop an efficacy-pharmacodynamic (PD) model for tumor suppression. We adopted drug level (e.g., Area-Under-the-Curve or AUC in central compartment of the PK model) as surrogate endpoint for efficacy, assuming direct proportionality between exposure and efficacy. This is contrasted to neutropenia based on the following parameters: (i) toxicity event (i.e., occurrence of any neutropenic event), (ii) seven-day toxicity event (i.e., neutropenic event lasting for at least 7 consecutive days), (iii) recovery from neutropenia (i.e., recovery to Grade 1, meaning at least one ANC measure above Grade 2 threshold after a toxicity event), (iv) time to recover (i.e., time between first toxicity onset and first subsequent ANC above Grade 2). The toxicity events considered were neutropenia Grade 3 (ANC below 1E9 neutrophil/liter) and Grade 4 (ANC below 5E8 neutrophil/liter). The evaluation of seven-day neutropenia is preferred since Grade 4 neutropenia lasting 7 days or more is a dose limiting toxicity by protocol. Simulation analysis was limited to the first treatment cycle (28 days).

## Results

### Neutrophil life cycle model captures main stages of neutrophil maturation

The QSP workflow is shown in Figure 1A. It integrates three modules (i.e., PK, PD, neutrophil life cycle) and accessory operations (e.g., definition of virtual patients, model validation).

**Figure 1.**
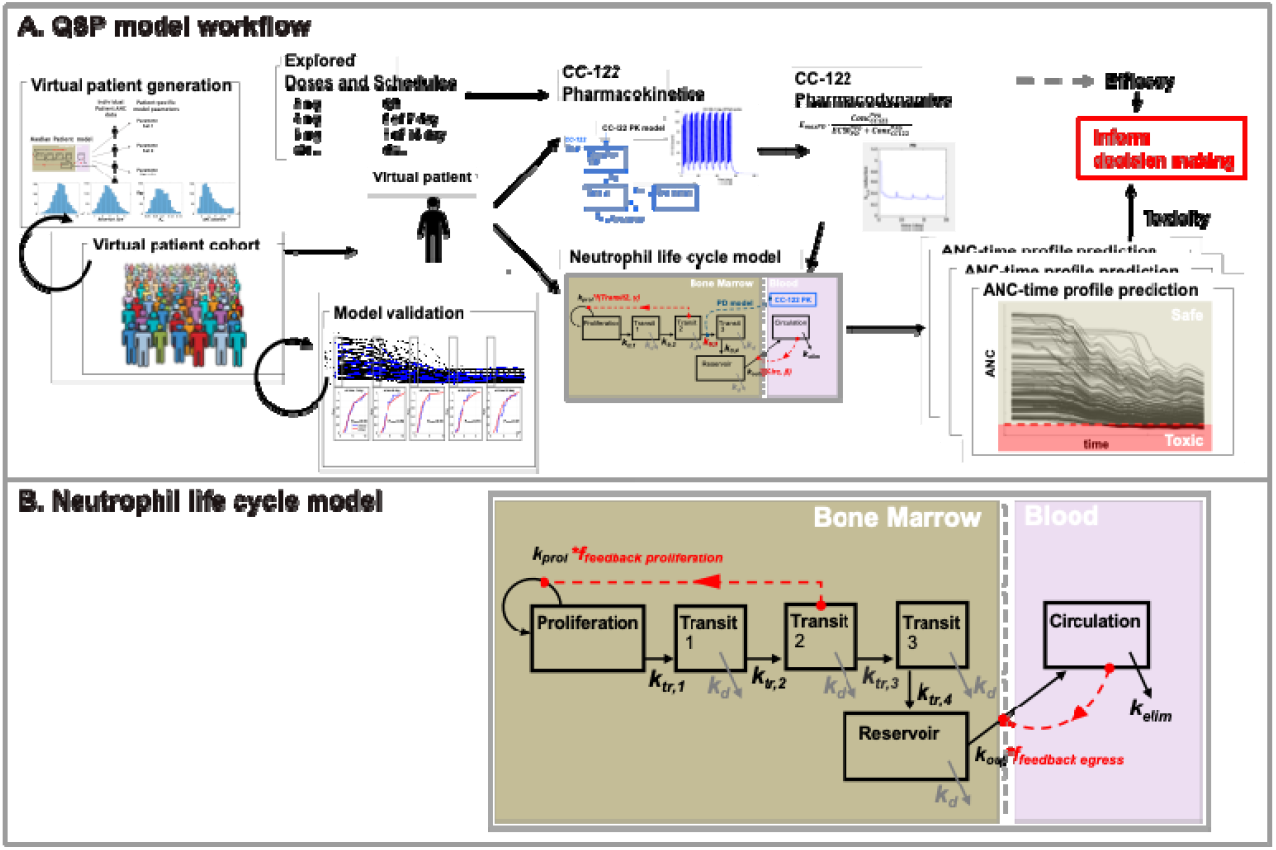
A: QSP model workflow. A virtual patient is represented as an appropriately parameterized model describing the neutrophil life cycle. This model can be solved to generate simulations of neutrophil counts in blood under homeostatic or avadomide-perturbed conditions. Avadomide effect is determined by the sequential evaluation of PK, PD, and PD-driven alteration of the neutrophil maturation. Model simulations iterated for a large cohort of virtual patients allow capturing the global pattern of neutropenia in the disease cohort under investigation. Finally, simulation results are postprocessed to compute toxicity endpoints of interest. B: compartmental structure of the neutrophil life cycle model. The proliferation pool represents committed proliferative neutrophil precursors. From a model idealization standpoint, these cells have specific characteristics: they can proliferate but not self-renew and can proceed to subsequent maturation stages, represented in the model as a sequence of transit compartments. These compartments (i.e., Transit1, Transit 2, and Transit 3) do not have a direct biological counterpart but here are intended to capture the fact that progressive maturation implies a time-delay, in line with previously published implementations of neutrophil maturation models. Once maturation is completed, cells are stored in a bone marrow Reservoir pool, awaiting egress into peripheral blood circulation. Circulation pool represents circulating neutrophils (i.e., level of neutrophils in blood, comparable to clinical ANC). Finally, circulating neutrophils are subjected to terminal elimination (cell death).

The neutrophil life cycle model (Figure 1B, Equations 1-8) describes neutrophil formation and maturation processes in bone marrow hematopoietic space, neutrophil egress from bone marrow to peripheral blood circulation, and neutrophil terminal death. The model consists in a proliferation pool (Proliferation), with proliferation rate *k*_*prol*_; a sequence of maturation stages (Transit 1, 2, 3) with sequential, first-order transfers and rate constants *k*_*tr,1*_, *k*_*tr,2*_, *k*_*tr,3*_, *k*_*tr,4*_; a reservoir pool (Reservoir) of mature neutrophils stored in bone marrow and final release to peripheral blood (Circulation). Bone marrow egress is controlled by the *k*_*out*_ rate constant. Finally, circulating neutrophils are subjected to terminal death based on *k*_*elim*_ rate, while maturing neutrophils undergo apoptosis based on *k*_*d*_ rate constant.

The model formulation was adapted to capture the specificity of the avadomide mechanism of action and to acknowledge the role of Ikaros upon neutrophil maturation. The *k*_*tr,3*_ expression was modified into a Michaelis-Menten based functional form (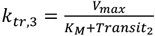, in Equations 3-4). The model includes two regulatory feedback mechanisms of neutrophil maturation under perturbed conditions: Feedback Proliferation (Equation 7) modulates the proliferation rate based on Transit 2 level and Feedback Egress (Equation 8) regulates egress of neutrophils from reservoir pool to peripheral blood. Both feedback mechanisms have a similar functional form, the exponents (γ and β) modulate the velocity of the control action. For full details of model formulation refer to Supplementary Materials 2.1.

### Avadomide PK and PD models

The avadomide PK is described by a two-compartment PK model. The avadomide PD model (Equation 9) determines the magnitude of neutrophil maturation block as a function of avadomide concentration (details in Supplementary Materials 2.2).

### Clinical trial data show high inter- and intra-disease cohort variability in longitudinal ANC patterns

We conducted a preliminary data analysis to explore patterns of longitudinal ANC profiles for the first treatment cycle (Figure 2) across and within disease cohorts and dosing groups. This analysis revealed a significant variability in the longitudinal ANC profiles that associated with both initial patient characteristics (e.g., baseline ANC measures from ~2E9 to 8E9 cell/liter, Figure 2A) and treatment dosing schedules (normalized nadir depth varies within the same disease cohort for different dosing schedules, Figure 2C). These results emphasize the need to generate disease-specific models and the importance of capturing patient variability within individual cohorts.

**Figure 2.**
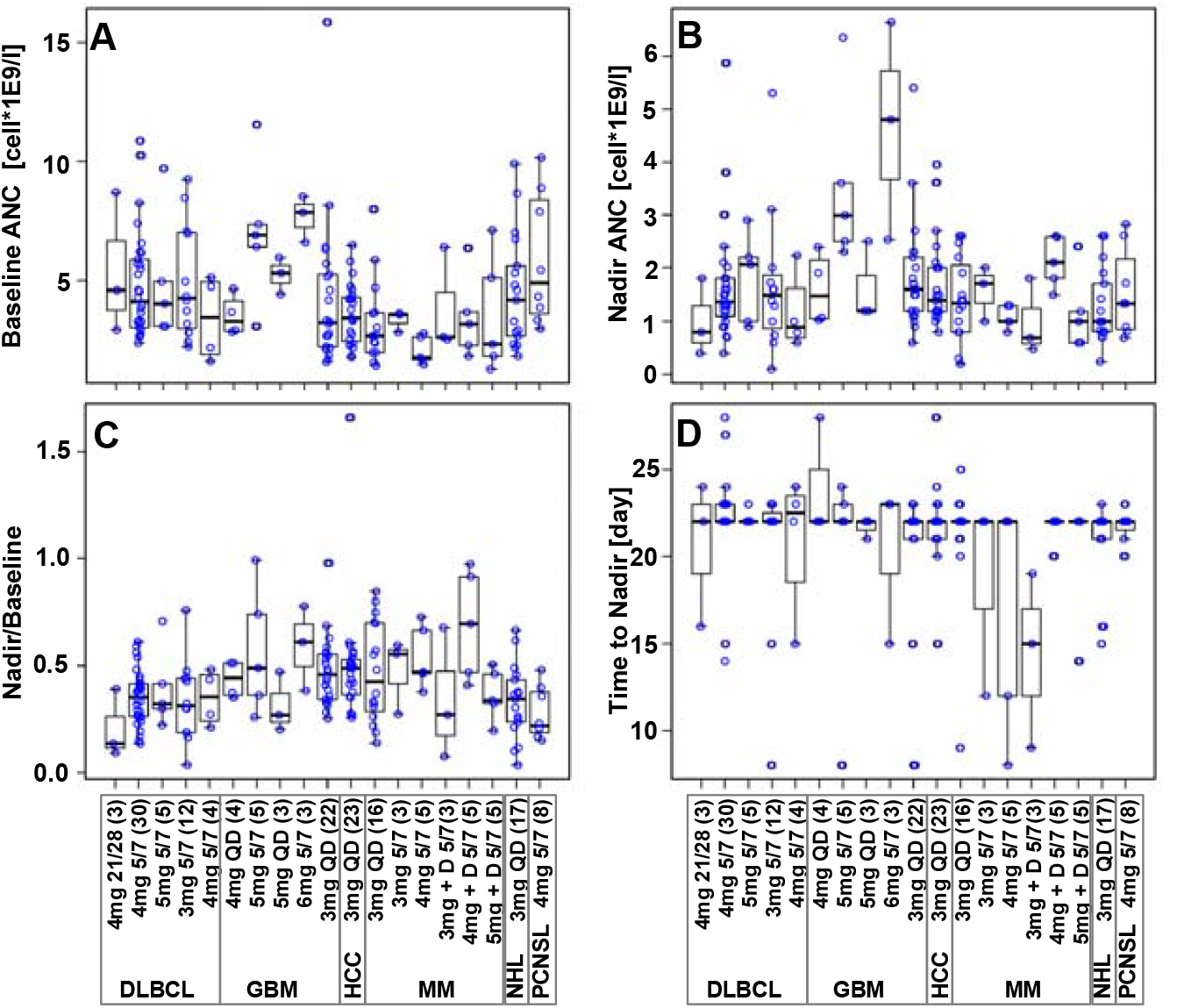
Boxplots of ANC patterns for avadomide-treated patients in multiple disease cohorts. Blue dots show data for individual patients. A: Average of available ANC measurements prior to treatment start; B: Lowest ANC measured within first treatment cycle; C: Nadir normalized to baseline; D: Time of nadir (typically day 22, however this result is conditioned by clinical sampling schedule, true value expected between days 16 and 28). Text boxes at the bottom indicate disease cohorts, specific doses and schedules, and number of patients in parenthesis. For MM cohort, “+D” label means avadomide + dexamethasone. NCT01421524 trial cohorts included patients with Glioblastoma (GBM), Multiple Myeloma (MM), Diffuse Large B-Cell Lymphoma (DLBCL), Hepatocellular Carcinoma (HCC) and Primary Central Nervous System Lymphoma (PCNSL). (References to related avadomide clinical trial data and data processing details in Supplementary Materials 1.3).

### Model parameterization explains disease cohort differences in ANC patterns

Model parameterization involved a combination of literature information, experimental observations, calculation, and regression. Because the neutrophil life cycle model (detailed in Supplementary Material 2.1) has a unidirectional and sequential transit compartment structure, most of the parameters can be calculated given one of these transit rates. We informed *k*_*elim*_ from literature and fixed *k*_*d*_ to a minor/negligible rate (as detailed below), and backward calculated *k*_*out*_, *k*_*tr,4*_, *k*_*tr,3*_, *k*_*tr,2*_, *k*_*tr,1*_, *k*_*prol*_ under the assumption of homeostasis (i.e., cell count remain constant in all compartments). Calculation details are shown in Table I.

**Table I.**
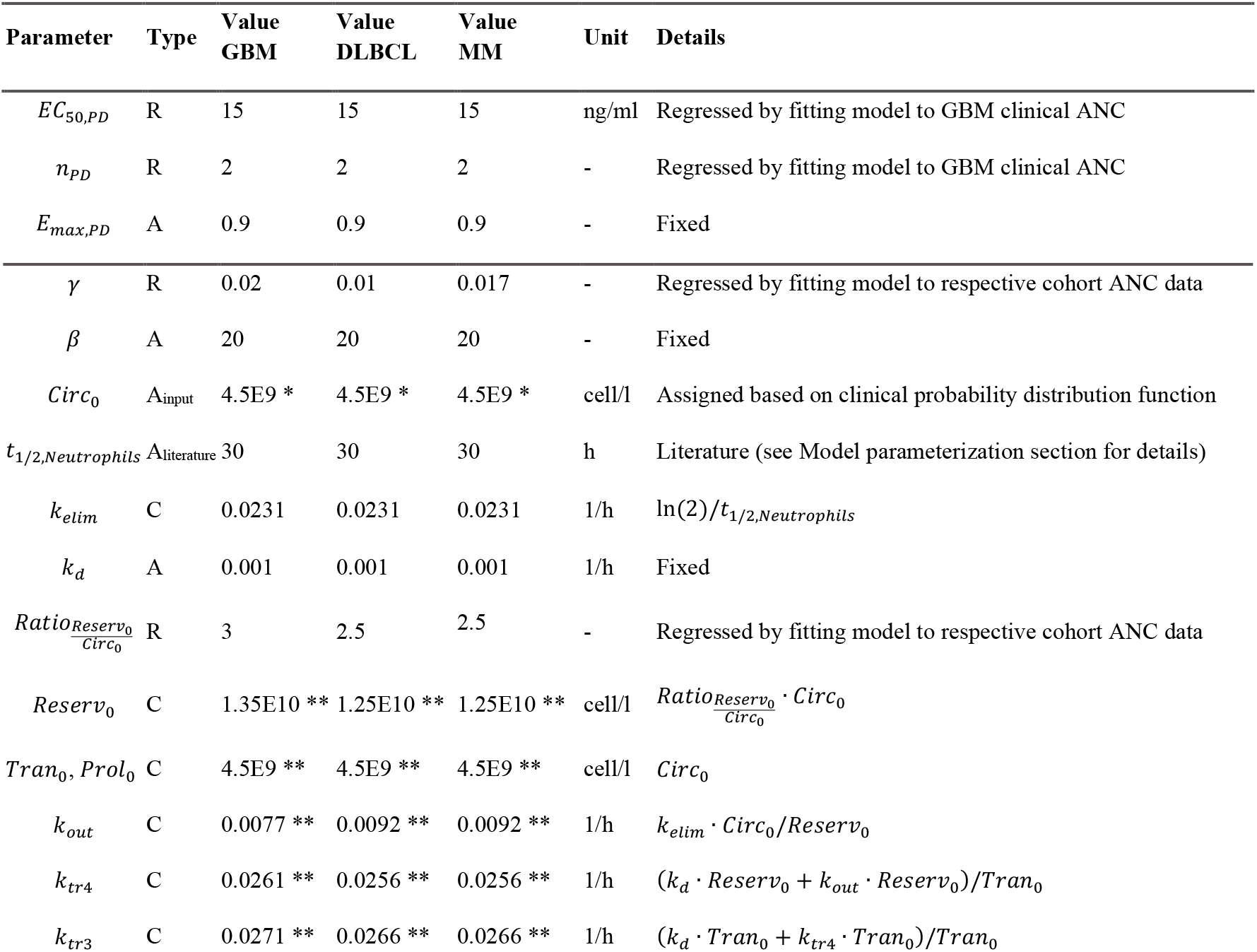
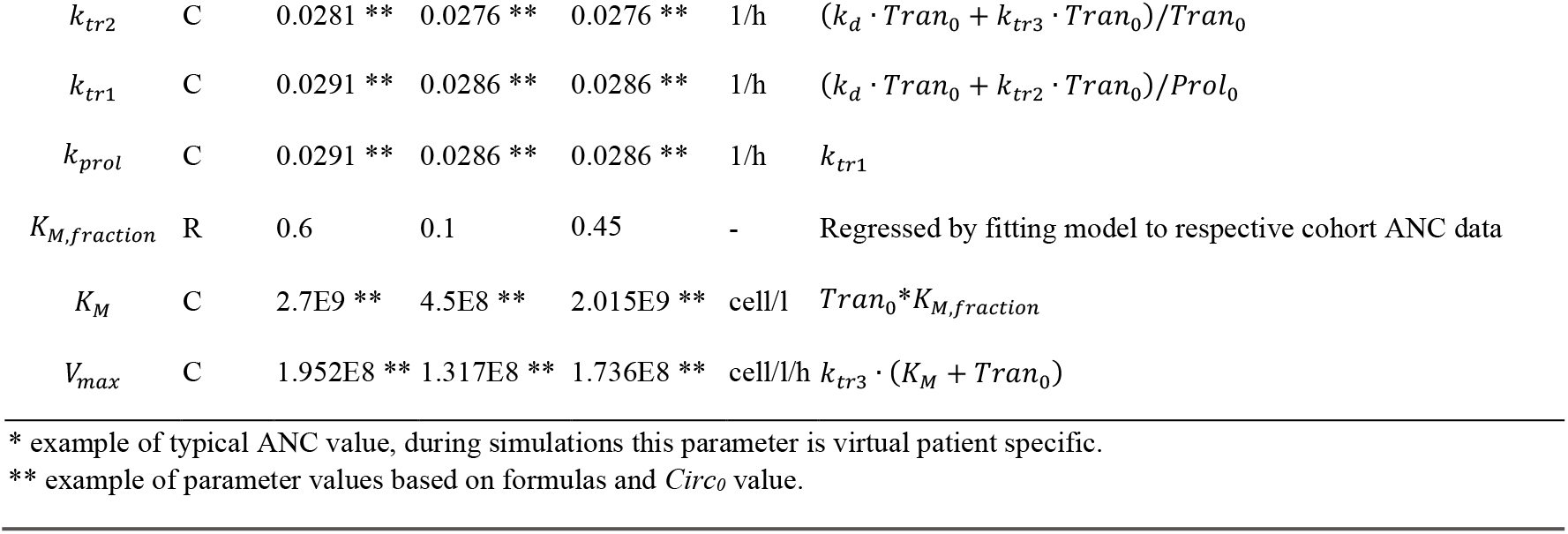
Model parameters for avadomide PD and neutrophil life cycle (median) model for GBM, DLBCL, and MM. Type column refers to parameter assignment: A=assigned from literature or fixed arbitrarily; C=computed based on equation reported in the Details column; R=regressed.

The half-life of circulating neutrophils in humans is subject of discussion. Several publications report contrasting data (21,37–39), proposing that half-life could range from a few hours to several days. Difficulty in measuring this parameter depends mostly on the cell-labeling system adopted and to the fact that neutrophils can relocate to marginated sites thereby affecting apparent circulating half-life estimates. Furthermore, neutrophil life-span can change under non-homeostatic conditions (39). In particular, Dale *et al*. (40) reported that under neutropenic state, neutrophil life span doubles (t_1/2_ = 9.6 h control vs 20.3 h neutropenia state). Given this knowledge and because the majority of papers report half-life ranging from 4 to 18 h (39), with a recent report measuring 3.8 days (41), we choose a typical value of 15 h and we double it to 30 h in agreement with enhanced life-span for neutropenia disease state. Finally, because all transit parameters are related, the choice of a different t_1/2_ within this range would not lead to significant changes in model outputs.

For initial cell count in the model compartments, because it was not possible to determine neutrophil cell concentration in the human hematopoietic tissue *in vivo*, we adopted the same approach of Friberg et al. 2002 (16) and fixed the initial cell level in all compartments (excluding the Reservoir component) to the initial neutrophil concentration in blood.

Remaining parameters were regressed or fixed to constant values. Regressed parameters include: the exponent of the Feedback Proliferation function (*γ*); the initial cell level in the reservoir pool (expressed as the ratio of cell level in the reservoir pool divided by cell level in Circulation, or *Ratio*_*Reserv0/Circ0*_), and *K*_*M*_ (in the following expressed as fraction of the initial cell level in Transit2 compartment, or *K*_*M, fraction*_). These parameters allow modulation of neutropenia patterns in different disease cohorts (e.g., GBM or DLBCL patients) or across individual patients and are discussed below. Fixed parameters are *k*_*d*_ and *β. k*_*d*_ was introduced above as a maturing cell death rate. The *in vitro* maturation assay showed that avadomide induces a reversible maturation block with no significant change in cell viability. However, apoptosis of maturing cells is a biologically recognized process and it is possible to speculate that *in vivo* neutrophils undergoing long term maturation block may experience enhanced apoptosis. Based on this, we included this process in the model with an arbitrarily assigned small rate (i.e., 0.001 h^−1^ or ~ 4% of *k*_*tr*_ maturation rates departing from the same compartments). The parameter *β* controls egress rate from the bone marrow reservoir pool. The biological mechanism controlling neutrophil egress from bone marrow is complex and only partially understood (42). We fixed *β* to a high value based on the clinical observation that, even in presence of avadomide block, circulating ANC was stably maintained at baseline level for several days despite compromised bone marrow maturation, suggesting that the egress of mature neutrophils from bone marrow is sustained and prompt.

The model was initially fitted to data from GBM patients. Those patients did not receive previous lines of bone marrow depleting treatments and therefore represent the closest match to a healthy bone marrow condition before avadomide treatment. The model was fit simultaneously to all GBM dose groups in order to regress a single parameter set representative of the GBM patient population (Figure 3A). At this step, five parameters were fitted. Three of those parameters are disease-group specific: *γ, Ratio*_*Reserv0/Circ0*_, *K*_*M, fraction*_, and two are PD specific: *EC*_*50,PD*_ and *n*_*PD*_. Once regressed, PD parameters are kept constant for any other avadomide simulation/fit under the assumption that drug effect is reproducible across the disease cohorts. The three disease-group specific parameters are instead re-fitted per disease group, because these parameters are representative for the bone marrow state and thus change across disease cohorts.

**Figure 3.**
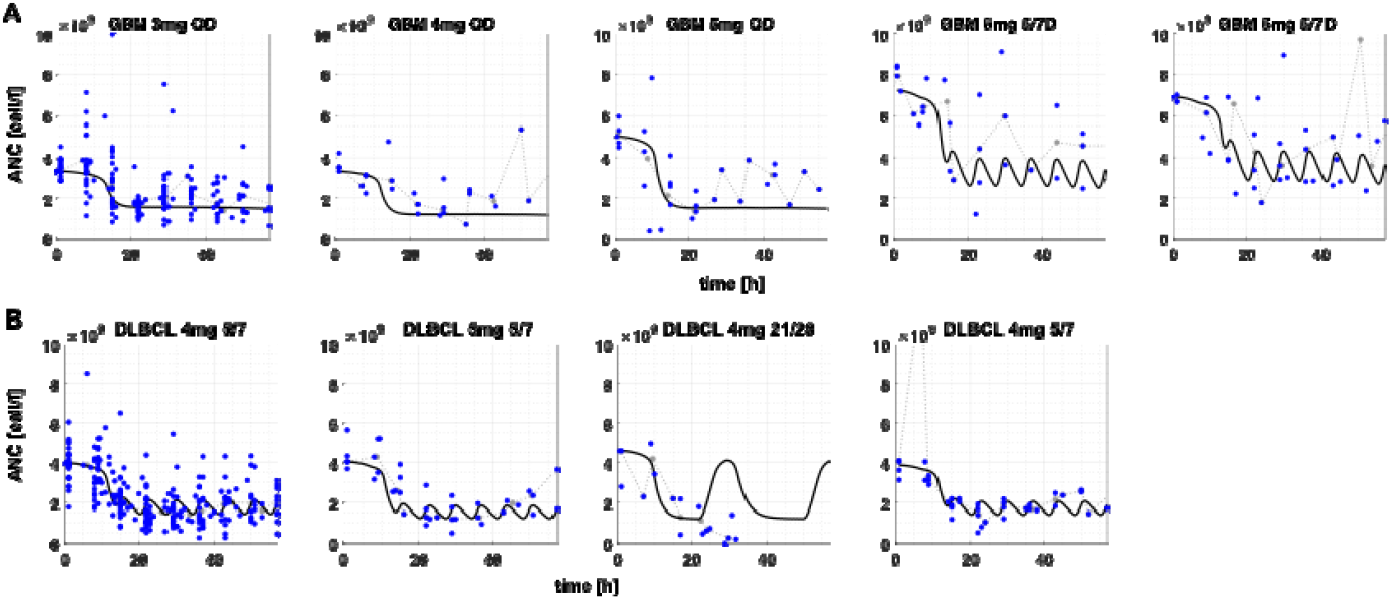
A: Model best-fit to ANC data for all GBM dose groups; B: Model best-fit to ANC data for multiple DLBCL dose groups. Legend: Black-solid line: model fit; gray-dotted line: clinical ANC median profile; blue dots: individual (processed) clinical ANC. Schedules: QD=daily dosing; 5/7=5-days on, 2days-off; 21/28=21-days on, 7days-off.

For model fit to the DLBCL median profiles (i.e., gray dotted lines in Figure 3B), the parameters *γ, Ratio*_*Reserv0/Circ0*_, *K*_*M, fraction*_ were refitted starting from the GBM estimate as initial guess. This operation served multiple purposes: (i) determine typical parameter values of DLBCL patients, (ii) explore whether parameter value differences between GBM and DLBCL could explain biological differences between the two patient groups, and (iii) determine initial parameter estimates for the subsequent step of patient-specific model fits.

Figure 3B shows a model fit to median DLBCL ANC data and Table I compares fitted parameter values for GBM vs DLBCL. It can be observed that parameters representing size of mature neutrophil reservoir pool in bone marrow (i.e., Ratio_Reserv0/Circ0_), extent of proliferative response to avadomide maturation block (i.e., γ), and idiosyncratic capacity to contrast maturation block (i.e., *K*_*M, fraction*_) are reduced in DLBCL compared to GBM.

### Virtual patient cohort

The following four model parameters allow for characterization of individual patients: (i) ANC level at baseline, (ii) size of the neutrophil reservoir pool in the bone marrow, (iii) *K*_*M*_ parameter in the Michaelis-Menten formulation of *k*_*tr,3*_, and (iv) γ exponent in the Feedback Proliferation function. Briefly, the ANC level at baseline is the neutrophil count in blood before treatment start. The size of the neutrophil reservoir pool represents individual initial level of mature neutrophils stored in bone marrow at treatment start (it influences the time needed before a drop in circulating ANC is observed). The *K*_*M*_ parameter regulates changes to neutrophil transfer from Transit 2 to Transit 3 when Transit 2 cell level deviates from its homeostatic value. The γ exponent controls the magnitude of proliferative response to the avadomide-induced perturbation of neutrophil maturation.

Starting from the DLBCL reference parameter set, the model was re-fitted to individual ANC profiles in the DLBCL cohort, thereby generating a set of values for each parameter. Because not all parameter value distributions are normal, we kept the parameter empirical distributions as they are (i.e., without replacing them with parametric models) and adopted kernel density estimation to estimate the probability density function (Figure 4A).

**Figure 4.**
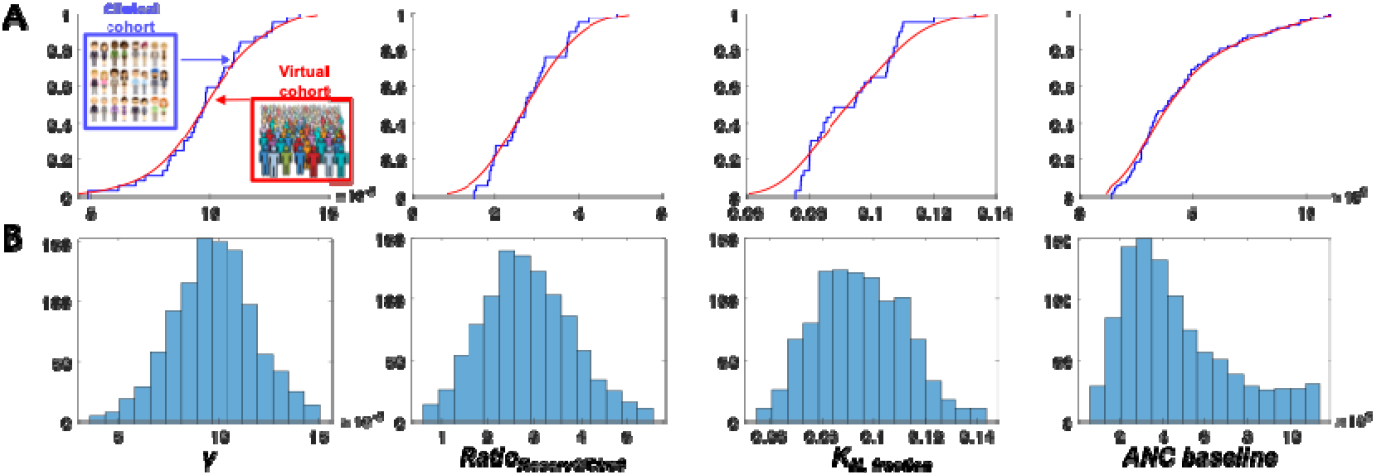
Virtual cohort generation. A: cumulative empirical distributions for DLBCL fitted-parameter values (blue) vs probability density function estimates (red). B: histograms of final parameter value distributions for 1000 virtual patients.

Finally, virtual patients were created by independent random sampling from the parameter value probability distribution functions (parameter values are assumed independent, meaning that there is no conditional probability for parameter values given the value of other parameters). The virtual cohorts generated for this analysis included 1,000 virtual patients (Figure 4B).

### Model identifiability and global sensitivity analyses

The model was tested for identifiability considering the three individualized parameters (*γ, K*_*M, fraction*_, *Ratio*_*Reserv0/Circ0*_) and specifying that observations are only available for Circulation compartment. *K*_*M, fraction*_ and *Ratio*_*Reserv0/Circ0*_ are globally structurally identifiable, while *γ* is locally identifiable.

We used GSA to rank parameters by importance in determining changes to the simulated ANC profile (full results in Supplementary Materials 2.4). GSA results support the choice of *γ* and *Ratio*_*Reserv0/Circ0*_ as individual parameters for the generation of the virtual patient population, while indicate that *K*_*M, fraction*_ is likely to contribute poorly toward differentiating virtual patients. For the present application, we acknowledge the minor role of this parameter, which could nonetheless be relevant for model application in the context of other indications and it is therefore kept in the virtual patient generation workflow.

### Virtual population of DLBCL patients reproduces clinically observed longitudinal ANC profiles

The virtual DLBCL patient population was validated by simulating the same treatment received by two clinical trial cohorts (avadomide 3 mg on a 5/7 and QD schedule, data not used to generate the virtual population) and then testing equivalence of the virtual and the clinical ANC distributions at selected times. Figure 5 shows how these distributions were found being equivalent at all tested times for the 3mg QD group and for 4 of 5 times for the 3mg 5/7 group.

**Figure 5.**
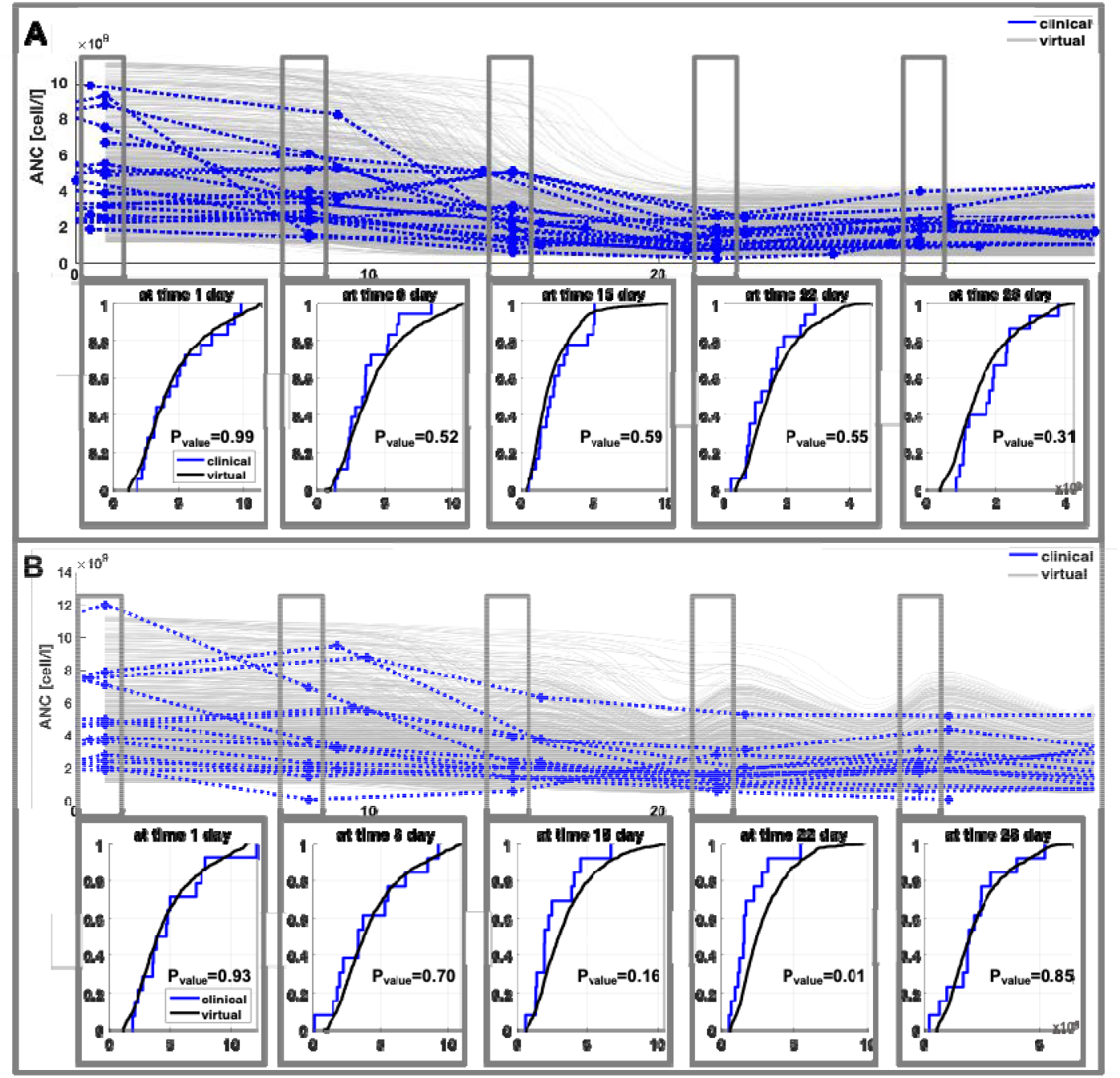
Model validation results. A: Avadomide 3 mg QD. Top: longitudinal ANC profiles, virtual cohort (1000 subjects) = gray-solid, clinical cohort (18 patients) = blue-dotted. Bottom: K-S test for equivalence of cumulative distribution profiles (with 5% significance level P_value_). B: Avadomide 3mg 5/7 day. Top: longitudinal ANC profiles, virtual cohort (1000 subjects) = gray-solid, clinical cohort (14 patients) = blue-dotted. Bottom: K-S test for equivalence of cumulative distribution profiles (with 5% significance level P_value_). Virtual and clinical ANC distributions were taken at day 1, 8, 16, 22, and 28 and compared using the two sample K-S test. Distribution equivalence rejected only for 3mg 5/7 at day 22 (i.e., equivalence verified at day 1, 8, 16, 28, but not at day 22).

### Model is applied to explore doses and schedules

Avadomide administration to the virtual DLBCL cohort (1000 virtual patients) was simulated for all combinations of 7 doses (i.e., 2, 3, 4, 5, 6, 7, 8 mg) and 6 schedules (i.e., 3/7, 5/7, 7/14, 14/28, 21/28, 28/28), totaling 42,000 simulations. Next, individual predictions of ANC profiles were processed to determine whether or not avadomide caused Grade 3 or 4 neutropenia, its duration, the recovery, and the time to recover. Collective analysis determined the percentage of patients expected to experience toxicity and possibly recover from it within the first drug administration cycle. Here we report a selection of representative results, full results available in Supplementary Materials 2.5.

Figure shows the longitudinal ANC profiles for the same virtual cohort receiving 6 mg of avadomide on the 5/7 or 21/28 schedule. In terms of exposure, the two schedules allow similar total dosing and PK exposure over the first cycle (20 doses and 1417 ng/ml*h AUC_cycle1_ vs 21 doses and 1515 ng/ml*h AUC_cycle1_, for schedules 5/7 and 21/28 respectively). Simulations show that until exhaustion of the reservoir pool, the ANC level remains stable, whereas at later time points (typically after day 10 post administration) ANC start dropping towards neutropenic levels. The schedule 5/7 shows that ANC nadir is reached for most virtual patients by day 21 with very few Grade 4 events, typically of short duration (~3 days). Virtual patients on the 21/28 schedule are shown to reach neutrophil count very proximal to absolute nadir by day 15 with a higher portion of patients experiencing Grade 4 neutropenia. Furthermore, ANC profiles for the 21/28 schedule are maintained proximal to nadir for several days, however the 7-day dose interruption enable a substantial recovery to level proximal to baseline. In both scenarios, ANC longitudinal profiles are tightly bound to dosing schedule.

Table II shows incidence of high-grade neutropenia and recovery for (i) different schedules at the same dose (4 mg) and for (ii) same schedule at different doses (5/7, 2 to 8 mg).

**Table II.** Summary of simulation results for different avadomide dosing schedules in virtual DLBCL cohort. A: multiple schedules for an avadomide 4 mg dose. B: different doses of avadomide given by a 5/7 schedule. Gr3 (Grade 3) and Gr4 (Grade 4) single indicate percentage of virtual patients experiencing at least one event of neutrophil level below the respective toxic threshold. Gr3 and Gr4 7 days indicate the percentage of virtual patients experiencing extended and uninterrupted Grade 3 and 4 toxicity, respectively, for at least 7 consecutive days. Recovered Gr3 to above Gr2 and Gr4 to above Gr2 indicate the percentage of patients that recovered to Grade 1 (i.e., above Grade 2). Analysis is limited to the first treatment cycle.

|  | Gr3<br>single<br>[%] | Gr4<br>single<br>[%] | Gr3<br>7 days<br>[%] | Gr4<br>7 days<br>[%] | Recovered<br>Gr3 to<br>above Gr2<br>[%] | Recovered<br>Gr4 to<br>above Gr2<br>[%] | Mean time<br>to recover<br>from<br>Gr3 to<br>above Gr2<br>[day] | Mean time<br>to recover<br>from<br>Gr4 to<br>above Gr2<br>[day] | AUC<br>[ng/ml*h] | C<sub>max</sub><br>[ng/ml] |
| --- | --- | --- | --- | --- | --- | --- | --- | --- | --- | --- |
| **Schedule** | **A. Multiple schedules for avadomide 4 mg dose** |  |  |  |  |  |  |  |  |  |
| 3/7 | 5.3 | 0 | 1 | 0 | 0 | 0 |  |  | 571 | 91 |
| 5/7 | 25.9 | 3.9 | 8.9 | 0 | 0 | 0 |  |  | 945 | 96 |
| 7/14 | 19 | 2.6 | 3.3 | 0 | 12.5 | 0 | 4.67 |  | 672 | 96 |
| 14/28 | 33.7 | 5.9 | 9 | 0.5 | 28 | 1.4 | 6.26 | 9.51 | 676 | 98 |
| 21/28 | 45.4 | 9.2 | 36.6 | 6.8 | 38.5 | 2.4 | 11.24 | 11.71 | 1010 | 98 |
| 28/28 | 45.9 | 9.6 | 45.6 | 9.1 | 0 | 0 |  |  | 1303 | 98 |
| **Dose [mg]** | **B. Multiple doses for avadomide on 5/7 administration schedule** |  |  |  |  |  |  |  |  |  |
| 2 | 5.5 | 0 | 2.7 | 0 | 0 | 0 |  |  | 472 | 48 |
| 3 | 13.5 | 0.2 | 5.4 | 0 | 0 | 0 |  |  | 709 | 72 |
| 4 | 25.9 | 3.9 | 8.9 | 0 | 0 | 0 |  |  | 945 | 96 |
| 5 | 36.7 | 6.5 | 13.2 | 0.2 | 1 | 0 | 2.69 |  | 1181 | 119 |
| 6 | 45.8 | 9.6 | 20.4 | 1.8 | 0.8 | 0 | 2.74 |  | 1417 | 143 |
| 7 | 53.9 | 12.4 | 27.3 | 4.1 | 0.5 | 0 | 2.43 |  | 1653 | 167 |
| 8 | 59.7 | 15.7 | 33.7 | 5.4 | 0 | 0 |  |  | 1889 | 191 |

Based on Table IIA, drug exposure (measured as AUC) increases with the total number of dosing days while C_max_ increases with the number of consecutive dosing days. For neutropenia, the incidences of both Grade 3 and 4 neutropenic events increase with consecutive dosing days, with the exceptions of 5/7 which shows slightly higher incidence than 7/14. In contrast the incidence is not directly dependent to the total dose received, as shown by the differences between 7/14 vs 14/28 or 5/7 vs 21/28. Interestingly, incidence of Grade 3 and 4 events is very similar for schedules 21/28 and 28/28. In contrast, this similarity is not found for neutropenia maintained for at least seven consecutive (7+) days, where we observe a substantial difference between schedules 21/28 and 28/28 which show incidence of 36.6%, and 45.6% (for Grade 3, 7+ days), respectively. For 28/28 single and 7+ day, neutropenia has same total incidence, while intermitted schedules show a reduction of 7+ neutropenic events compared to single events. In terms of recovery, all the intermittent schedules with at least 7 days of dose interruption show substantial recovery (i.e., 66% (12.5/19), 83% (28/33.7), and 84% (38.5/45.4) of virtual patients that experienced neutropenia Grade 3 recovered above Grade 2 for 7/14, 14/28, and 21/28, respectively). In contrast, no recovery was determined for 3/7 and 5/7 schedules. For schedules that allow recovery, the recovery time increases non-linearly with consecutive dosing days (i.e., 4.7, 6.3, and 11.2 days were necessary on average to recover from Grade 3 to above Grade 2 for schedules 7/14, 14/28, and 21/28, respectively).

Based on Table IIB, both AUC and C_max_, increase linearly with the dose. For neutropenia, the incidences of both Grade 3 and 4 neutropenic events increase less than proportionally with dose (rapid relative increase of neutropenia incidence at low doses and reduced relative increase at high doses). It is also observed that, on a 5/7 schedule, there is very little, or absent, recovery at all doses. For the very few patients that would recover from neutropenia, the recovery time is short and compatible with the dosing interruption interval.

Figure 7 shows a bar plot comparison of toxicity and recovery across schedules for two doses (4 or 6 mg), to complement the results proposed in Table II. Bars are schedule-specific and are ordered by increasing drug exposure. The higher the number of consecutive dosing days the higher the percentage of patients experiencing toxicity. This pattern is not verified for 5/7 vs 7/14 likely because of the combined effect of similar dosing days (5 vs 7 days) and the difference in the dosing holiday (2 vs 7 days). Recovery from Grade 3 is substantial (>80%) and very similar for 14/28 and 21/28 and increases with dose for schedules 7/14 and 14/28, but not for 21/28. Increase in dose from 4 to 6 mg associates with higher recovery from Grade 4. Schedule 5/7 shows some lower toxicity compared to other schedules but offers little or no recovery.

**Figure 6.**
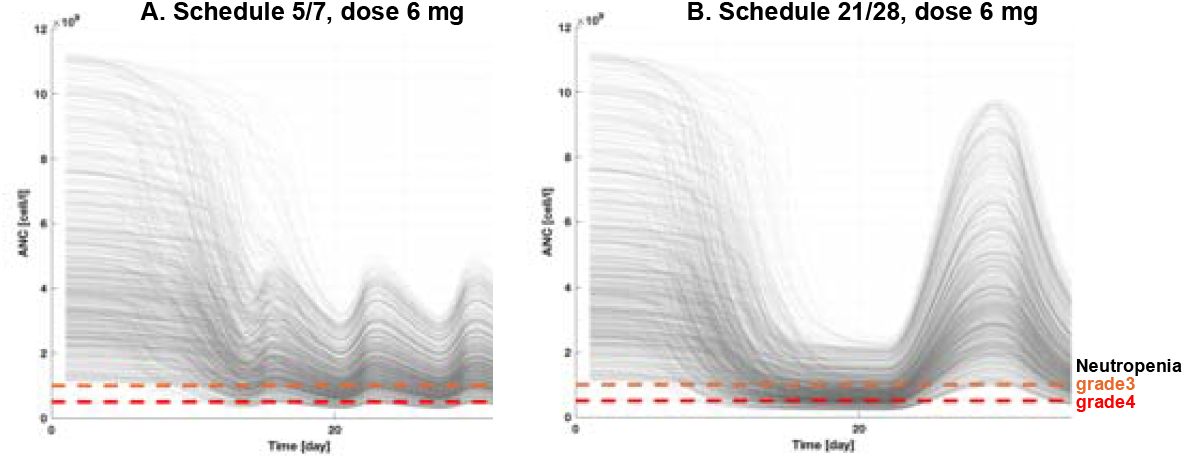
Simulation of the same 1000 virtual patients for avadomide 6 mg on a 5/7 (A) or 21/28 (B) schedule. Neutropenia Grade 3 (orange) and 4(red) are represented as horizontal dashed lines. The ANC baseline distribution (i.e., ANC at t=0) is the same because the same virtual patients are simulated for both dosing schedules. The two schedules enable very similar PK exposure over the first treatment cycle; however, the neutropenia pattern is quite different: schedule 21/28 shows deeper ANC drop and protracted toxicity, followed by strong recovery once the treatment is interrupted. In contrast, schedule 5/7 offers a mitigated incidence of high-grade toxicity, with only limited recovery during dose interruption.

**Figure 7.**
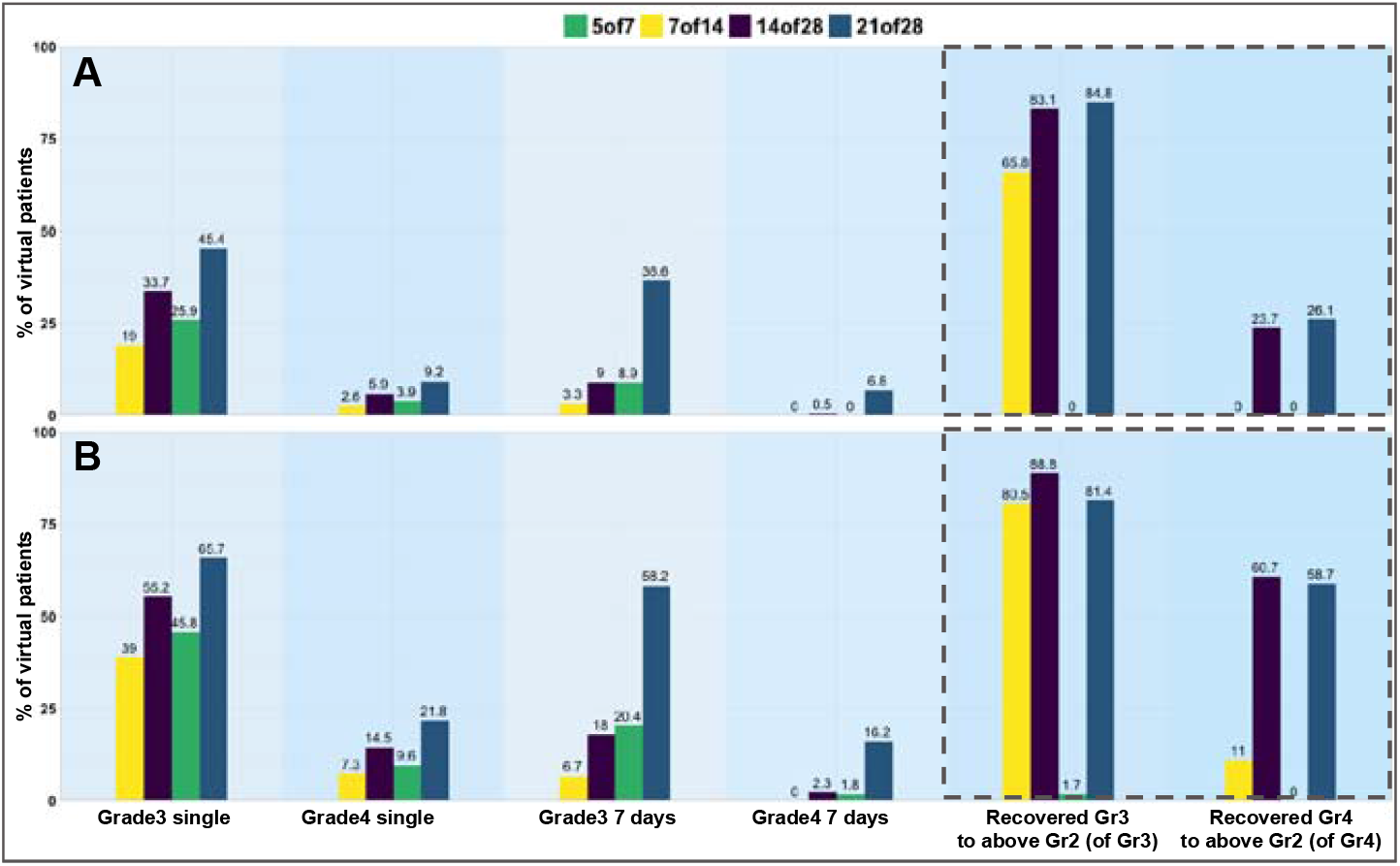
Bar plot analysis for toxicity and recovery for different schedules at 4 mg (A) and 6 mg (B). Grade 3 and 4 single indicate percentage of virtual patients experiencing at least one event of neutrophil level below the respective toxic threshold. Grade 3 and 4 7 days indicate percentage of virtual patients experiencing an extended and uninterrupted toxicity for at least 7 days. Recovery Gr3 to above Gr2 and Gr4 to above Gr2 indicate the percentage of patients that recovered to Grade 1 (i.e., above Grade 2) relative to the patients that experienced toxicity. This analysis is limited to first treatment cycle.

Figure 8 shows the time of nadir for five different schedules. Schedule 5/7 shows bimodal time of nadir with ~9% of patients having nadir at day 20 and ~91% at day 27. Schedule 7/14 and 21/28 show nadir at day 21, consistently with the start of the latest dosing holiday for cycle 1. Schedule 14/28 shows nadir in the interval of day 15 to 17. Finally, daily dosing (schedule 28/28) results in progressive increase of the virtual patients having ANC nadir in the interval of day 21 to day 28.

**Figure 8.**
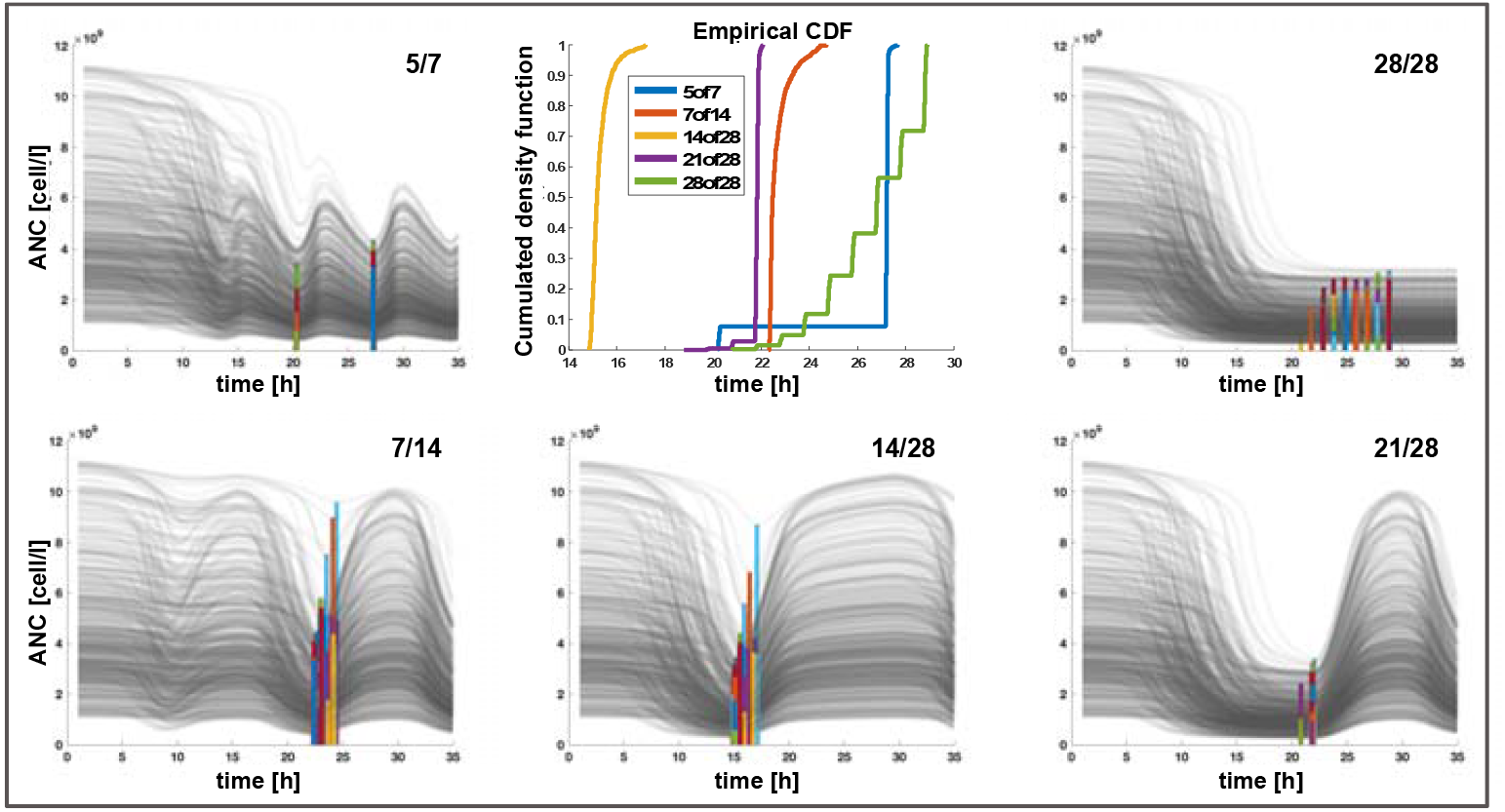
Time of nadir across schedules. Central top panel shows the empirical cumulative distributions of the time of occurrence of nadir for different schedules. Surrounding plots offer a visual justification for the observed nadir-time pattern. These plots show longitudinal ANC profile for 500 virtual patients with graphical visualization of individual nadirs by vertical-colored bars. Bar heigh depends on the individual ANC at nadir.

## Discussion

In this paper we have presented a QSP model for avadomide induced neutropenia. We applied this model to virtually explore the pattern and the incidence of neutropenia across dosing schedule scenarios in a DLBCL patient population treated with avadomide. Model development followed good practice standards as described in Bai et al. 2019 (23).

The neutrophil life cycle model developed describes neutrophil maturation and transit stages from bone marrow to peripheral blood and captures the avadomide-specific mechanism of induction of neutropenia. Since this mechanism is different from chemotherapy-induced neutropenia, published models (such as the Friberg model (16)) could not be applied to address needs of our study. A major difference of our model compared to the Friberg model (16) is that proliferation rate is not controlled by ANC level changes compared to baseline in peripheral blood. That mechanistic implementation was not well-suited to description of the CELMoD-driven neutrophil maturation block, and upon testing produced indefinite accumulation of neutrophils at the maturation blocked stage and excessive proliferation (because during maturation block, proliferation would be continuously stimulated by the sub-baseline ANC level). Additionally, a first order modeling of the cell transit through maturation stages is not suitable for CELMoD-like maturation block. For example, the first order based transit (i.e., rate constant*cell level in upstream compartment) in presence of CELMoD-depressed maturation rate constant results in accumulation of cells at the affected maturation stage, which eventually would mathematically compensate for rate constant reduction and ultimately cause net flow to overcome the maturation block. Accordingly, we adopted a Michaelis-Menten like function for Transit stage 2 which allowed an asymptotic behavior of the flow out of Transit 2 despite an increase in accumulated maturing neutrophils.

In terms of the workflow, the clinically observed variability of ANC supported extending model simulation from a single median virtual patient to a virtual patient population. The DLBCL virtual cohort utilized in our simulations was validated comparing the cumulated distributions of the clinical and the virtual cohorts ANC at selected time points. This approach allowed for both qualitative and quantitative evaluation of equivalence of the two empirical cumulated distributions. An alternative and commonly adopted approach, like the visual predictive check, is conceptually similar in terms of comparing virtual vs clinical distributions, but it is more qualitative in nature.

The heterogeneity of the virtual population is observable in the simulated ANC profiles in terms of initial baseline, neutrophil reservoir pool size (ANC starts dropping from baseline level at different times), and idiosyncratic variability in response to maturation block (visible as overlapping profile in the recovery time interval). A limitation of the current implementation is that population PK was not included, as that would improve significantly the representation of the variability across the virtual population.

Model utility was demonstrated by simulating avadomide administration to a virtual DLBCL cohort. Since it was not possible to develop an avadomide efficacy module in absence of specific biomarkers or tumor suppression data, the drug exposure (i.e., AUC in central PK model compartment) is considered as a surrogate efficacy endpoint and here it is used as a reference to contrast schedule toxicity.

Simulation results address different aspects of neutropenia pattern modulation by choice of dosing schedule. Frequent dosing (i.e., schedules 28/28 and 5/7) produce high systemic exposure along with the highest incidence of neutropenia, compared to other schedules at same dose. It is also shown that two-day dosing holiday on the 5/7 schedule is sufficient to reduce significantly the total incidence of neutropenia in the virtual population (e.g., at the 4 mg dose, the schedule 5/7 compared to 28/28 gives ~28% less exposure, but it lowers incidence of neutropenia Grade 3 by ~44%). However, two-day holiday does not allow measurable recovery from high-grade neutropenia. This suggests that for avadomide in DLBCL patients a longer dosing holiday should be considered in case a more substantial recovery is desired. For example, compared to 5/7 and 28/28, all other tested schedules with measurable incidence of neutropenia enable substantial recovery (Figure 7). It is noted that the exploration of neutrophil recovery rate during dosing holiday is only possible with model-based tools since trial patients are typically undergoing sequential cycles of treatment and receive concomitant medications for the mitigation of neutropenia (such as G-CSF).

Regarding the analysis of prolonged high-grade neutropenia lasting at least seven consecutive days (7+ day), among those schedules allowing dosing interruption (excluding 28/28), schedule 21/28 results in higher incidence of prolonged neutropenia, coherently with the 21-day continuous dosing not allowing for intermittent recovery. The schedule 5/7, despite some mitigation enabled by the two days of dosing interruption, produces a 7+ day neutropenia comparable to schedule 14/28. Schedule 7/14 shows the best performance in terms of minimizing 7+ day toxicity at dose level 4 to 6 mg. Further, results show that under continued dosing, the maximal neutropenia would be reached by day 21 (or a few days earlier), since the total incidence of high-grade neutropenia is nearly equivalent for schedule 21/28 and 28/28 (Table II).

Finally, the model enables predictions of the time at which the most severe neutropenia is reached (i.e. ANC nadir, Figure 8), showing that nadir time is primarily controlled by the schedule of choice, rather than the dose level.

Collectively, these model-based results show that the choice of dose and schedule offers a powerful handle to modulate the neutropenia in terms of absolute incidence in the patient population, as well as the time of ANC nadir, duration of neutropenic state, and extent of recovery. These results demonstrate the model potential applicability as a support tool to inform decision making in the clinic. Simulation results should be interpreted in the light of clinical protocol definitions for dose limiting toxicity and maximum tolerated dose as well as efficacy considerations.

## Conclusions

Neutropenia is a major treatment-emergent and dose-limiting toxicity in trial patients treated with avadomide. Intermittent dosing is an option to manage this toxicity and different combinations of dose and schedule enable controlling the toxicity-efficacy tradeoff. Here we presented a QSP model for avadomide-induced neutropenia, which includes a mechanistic model of neutrophil life cycle combined with avadomide PK and PD. The complete workflow allowed capturing the disease cohort variability and enabled performing simulations for several dosing schedule scenarios, aiming at screening options that would minimize neutropenia while enhancing drug exposure.

This model is the first developed specifically for neutropenia caused by block in neutrophil maturation and is validated on clinical data. We anticipate further opportunities to apply, develop and demonstrate the relevance of this model given potential use of avadomide and other CELMoD compounds either as single agents or in combination to treat a range of indications.

## Acknowledgments

Yan Li (BMS) provided avadomide PK model. Fan Wu and Manisha Lamba (BMS) provided relevant feedback on model applicability and interpretation of results. Carla Guarinos, Alicia Benitez, Maria Dolores Jimenez, Estela G. Torano (BMS/CITRE) conducted *in vitro* experimental activity and provided relevant feedback on neutrophil maturation assay.

## Conflict of interest/disclosure

R.A.A., M.P., S.C., D.W.P., S.K., M.M., M.W.B.T., R.L., A.V.R declare employment at Bristol Myers Squibb.

M.P., S.C., D.W.P., S.K., M.M., M.W.B.T., R.L., C.C.S., A.V.R declare equity ownership in Bristol Myers Squibb.

## Contributions

R.A.A., C.C.S., and A.V.R. designed the research. R.A.A. and C.C.S. performed data processing and formal analyses in consultation with all other authors. M.P., D.W.P., S.K., and S.C. provided clinical and translational insights. M.M. contributed to model design. M.W.B.T., R.L. provided support for project development and revised the manuscript. A.V.R. supervised the project and provided data analysis and modeling insights. R.A.A. and C.C.S. wrote the manuscript. All authors edited the final version of the manuscript.

## Equations

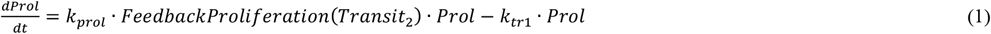

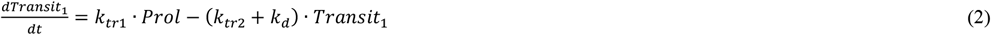

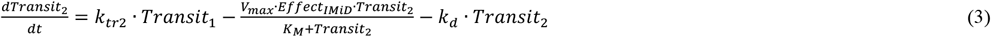

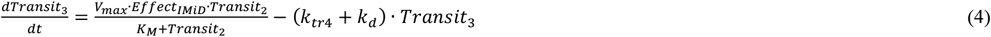

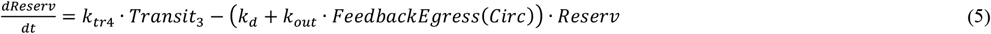

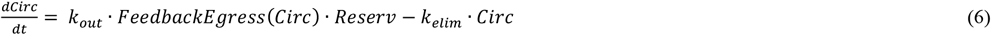

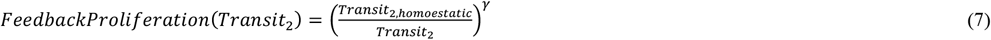

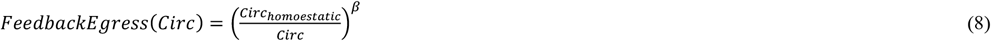

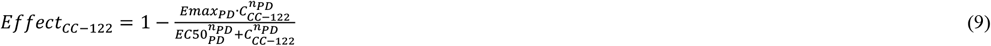

