## Supplementary Materials for "Quantitative systems pharmacology modeling of avadomide-induced neutropenia enables virtual clinical dose and schedule finding studies"

**Title:**

**Authors:**
Roberto A. Abbiati[1*], Michael Pourdehnad[2], Soraya Carrancio[3], Daniel W. Pierce[3], Shailaja Kasibhatla[3], Mark McConnell[4], Matthew W. B. Trotter[1], Remco Loos[1], Cristina C. Santini[5], Alexander V. Ratushny[4*]
**Institutions:**
[1]Bristol Myers Squibb, Center for Innovation and Translational Research Europe (CITRE), Seville, Spain
[2]Bristol Myers Squibb, San Francisco, CA, USA
[3]Bristol Myers Squibb, San Diego, CA, USA
[4]Bristol Myers Squibb, Seattle, WA, USA
[5]Current affiliation: Roche Pharma Research and Early Development, Pharmaceutical Sciences, Roche Innovation Center, Basel, Switzerland (Affiliation at the time the work was conducted: Bristol Myers Squibb, Center for Innovation and Translational Research Europe (CITRE), Seville, Spain)

### Supplementary Materials

#### 1 Methods

##### 1.1 Global Sensitivity Analysis

We adopted a Monte Carlo based method for Global Sensitivity Analysis (GSA), as described in Cho et al. (2003) (7). In brief, (i) the test range for each parameter was assigned, (ii) multiple parameter sets were defined by randomizing parameter values within their selected range by using Latin hypercube sampling (*lhsdesign* function in Matlab R2020a), (iii) model was repeatedly simulated for each randomized parameter set, (iv) model output was computed and contrasted to the reference simulation output obtained with the reference parameter set, and (v) for each parameter two cumulative distribution functions representing the simulations that produced output values above and below a selected reference output value were contrasted applying a Kolmogorov-Smirnov (K-S) test. Based on this method, the GSA parameter rank is determined by the test statistic, which consists of the maximum absolute difference between the two cumulative distribution functions.

#### 2 Results

##### 2.1 Neutrophil life cycle model details

The neutrophil life cycle model developed describes neutrophil formation and maturation processes in bone marrow hematopoietic space, neutrophil egress from bone marrow to peripheral blood circulation, and terminal death.

The model (Figure 1B, Equations 1-8) consists in a proliferation pool (Proliferation), with proliferation rate $k_{prol}$. A sequence of maturation stages (Transit 1,2,3) with sequential, first-order transfer and rate constants $k_{tr,1}$, $k_{tr,2}$, $k_{tr,3}$, $k_{tr,4}$. Mature neutrophils are stored in the reservoir pool (Reservoir) and eventually are released to peripheral blood (Circulation). Egress is controlled by the $k_{out}$ rate constant. Finally, circulating neutrophils are subjected to terminal death based on $k_{elim}$ rate. Maturing neutrophils undergo apoptosis based on $k_d$ rate constant.

The model formulation was adapted to capture the specificity of the avadomide mechanism of action and to acknowledge the role of Ikaros upon neutrophil maturation. Preclinical data showed that avadomide affects neutrophil maturation by inducing a reversible and incomplete block of cell maturation (1), which occurs primarily at the late maturation stages of neutrophil development (1) and does not affect proliferative neutrophil precursors (shown for lenalidomide (2)). Separate experiments suggested that this effect is related to an avadomide-mediated degradation of Ikaros. To implicitly implement these observations in the model, we modified the maturation process between Transit 2 and Transit 3 to represent a dual-agent interaction (i.e., maturing cell and Ikaros). $k_{tr,3}$ expression was modified into a Michaelis-Menten (M-M) based functional form ($k_{tr,3} = \frac{V_{max}}{K_M+Transit_2}$, in Equations 3-4). This empirical representation assumes that Ikaros level controls the net maturation/transit rate: an increasing cell level in Transit2 does not produce a linear increase in the net maturation/transit rate because Ikaros is the limiting agent (this is in contrast to classical first order representation), on the contrary a reduction of cell level in Transit2 reduces maturation rate.

The model includes two regulatory feedback mechanisms of neutrophil maturation under perturbed conditions: (i) Feedback Egress regulates egress of neutrophils from reservoir pool to peripheral circulation by increasing/reducing the egress rate based on neutrophil level in circulation to keep it stable at a set point level (i.e., homeostatic level). (ii) Feedback Proliferation modulates the proliferation rate based on Transit 2 level, to prevent excessive accumulation of maturing neutrophils in bone marrow. This is an empirical mechanism and does not intend to capture the real complexity of *in vivo* hematopoiesis regulation. When avadomide induces a reduction of neutrophil maturation (which was implemented as a $k_{tr,3}$ rate reduction between Transit stage 2 and 3), there is an accumulation of neutrophils at Transit 2. If unregulated, this process could lead to non-physiological cell levels in bone marrow, whilst in fact hematopoietic space has limited volume and can only accommodate a limited number of cells. It is acknowledged that this control system is specifically designed for our model idealization of the avadomide action and that hematopoiesis regulation mechanism can function differently from a biological perspective.

Mathematically, both feedback mechanisms have a similar functional form. Feedback Proliferation is formally described by Equation 7 and Feedback Egress by Equation 8. These contributions are activated when the cell level moves away from the homeostatic set point (i.e., ratio component of Equations 7 and 8 differs from 1). The exponents (γ and β) are parameters that modulate the velocity of the control action.

#### 2.2 PK and PD models

##### Avadomide PK

PK model structure: two-compartment PK model with first-order absorption to central compartment with absorption lag-time and first order elimination from central compartment.

PK model parameters:

| Parameter | Value | Unit |
|---|---|---|
| $k_{abs}$ | 5.5 | 1/h |
| $k_{12}$ | 0.0265 | 1/h |
| $k_{21}$ | 0.0195 | 1/h |
| $k_{el}$ | 0.0708 | 1/h |
| $Abs_{lag}$ | 0.406 | h |

Relevant publications: avadomide PK study (3).

Other related published PK models: pomalidomide (4).

##### Avadomide PD

*In vitro* experiments showed that the avadomide-induced block on neutrophil maturation is (i) incomplete (i.e., some degree of neutrophil maturation was preserved even at high drug concentrations), (ii) reversible (i.e., once avadomide was removed, mature neutrophils recovered to control level), and (iii) shows a concentration-dependent maturation block depth (1,5). Further, it appears that late- rather than early-maturation stages are most affected by the avadomide effect. Based on these observations, and by applying appropriate model simplifications, we imposed the PD effect at a single maturation stage between Transit stage 2 and 3. The concentration-effect relation is based on a Hill function (Equation 9), which generates a sigmoid-like curve. This model was adopted because preclinical data for avadomide (as well as lenalidomide) showed that a maximum maturation block is reached at increasing drug concentrations (1,2,5)

For parameters, $E_{max,PD}$ is the maximum maturation block, $n_{PD}$ is the Hill coefficient, and $EC50_{PD}$ is the half maximal effective concentration. The model mathematically describes the maturation rate reduction in comparison to its homeostatic value. The input to this PD model is avadomide concentration ($C_{CC\text{-}122}$) in patient's blood, which corresponds to avadomide in the central PK model compartment.

It was not possible to fit an independent PD model to data representing the concentration-maturation block relationship because of: (i) inability to measure "maturation block level" without measuring changes in the level of Ikaros; (ii) expected significant differences between neutrophil maturation *in vitro* vs *in vivo*; and (iii) the difficulty of translating drug concentration level from *in vitro* to *in vivo* due to absence of PK data at target site. For these reasons, it was not possible to fit all three PD parameters simultaneously, therefore $E_{max,PD}$ was fixed to a constant value (0.9, meaning 90% maximum maturation block) to acknowledge strong but incomplete block, and $EC_{50,PD}$ and $n_{PD}$ were determined by fitting the model to the patients' ANC data.

#### 2.3 Figure 3 data processing details

Prior to the model fit, a pre-processing of ANC data was performed to address the following issues: (i) the number of ANC samples and their collection times may not be exactly the same across patients (e.g., within the same interval of time, some patients had ANC measured a different number of times) and (ii) patients may have received concomitant medications (CM), which may have altered avadomide-monotherapy ANC level. As a result, when computing the averaged cohort ANC, patients with the greater/lesser amount of ANC measurements result in higher/lower weights towards the global average. For this reason, ANC data for individual patients were re-sampled by time window: individual ANC data within a 4-day window were replaced by their median values and the corresponding times by their average. This allowed equalization of sample number across patients while ensuring sampling uniformity over time. The time window of 4 days is similar to the clinical sampling schedule interval and was chosen as a compromise between having an overly small (excessive data granularity) or large (causing loss of resolution) window.

Finally, because the model does not describe CM effects formally (for this analysis, we considered only G-CSF related CM), ANC data measurements after first day of CM were removed from individual ANC profiles prior to further modeling analyses.

Related avadomide clinical trial data have been published previously (3,5,6)

#### 2.4 Global sensitivity Analysis

GSA was conducted based on the approach described in Methods and used the following settings: (i) parameter ($p_i$) randomization interval from $p_i/4$ to $p_i*4$; (ii) a 1000 parameter set generated via Latin hypercube randomization; (iii) the model outputs considered are neutrophil levels in the progenitor and circulation pools; (iv) the variation of the model output when simulation used the reference vs the randomized parameter sets was determined as the sum of normalized differences between the two output function values (this difference is evaluated for ANC in both progenitor and circulation compartments at 30 equally spaced time points from day 1 to day 28 and summed); (v) each output function value is compared to the global mean of all 1000 output function values to determine those simulations which gave higher vs lower output function value and thereby determining the two cumulative distribution functions used for the K-S test; (vi) since running these simulations for homeostatic condition would hinder the effects of some parameters, all simulations were run with an arbitrary avadomide administration (we chose the 3 mg and the 3/7 schedule because it is usually associated with an intermediate pattern of neutropenia severity) to explore a transient behavior of the neutrophil life cycle and its dependence on model parameter perturbations (explorative analysis with other schedules gave a comparable GSA ranking; results not shown).

Finally, it is noted that the choice of the model output can have a profound impact on GSA results. Along with the current GSA analysis shown here, alternative formulation for the model outputs have been tested and resulted in slightly different K-S statistics, but no significant change in final parameter ranking (results not shown).

GSA results are shown in Table S.I.

*Table S.I. Model parameter ranking based on the global sensitivity analysis. The two PD parameters $EC_{50,PD}$ and $n_{PD}$ ranked highly (1st and 3rd respectively) coherently with their role in modulating drug-induced effect on neutrophil level. $\gamma$ and $Ratio_{Reserv0/Circ0}$ ranked 2nd and 4th, respectively: $\gamma$ directly controls the proliferation response to the drug induced perturbation, while $Ratio_{Reserv0/Circ0}$ changes are directly observed in circulating neutrophil levels. Maturing neutrophil death rate ($k_d$, ranked 5th) has a measurable impact on neutrophil levels, making it a relevant parameter whose role in the model should be further explored. This result underscores the importance of accurate quantification of neutrophil death/apoptosis rates for mechanistic model predictions. Finally, $K_{M,fraction}$ and $\beta$ are low ranked (6th and 7th respectively).*

| **Parameter** | $EC_{50,PD}$ | $\gamma$ | $n_{PD}$ | $Ratio_{Reserv0/Circ0}$ | $k_d$ | $K_{M,fraction}$ | $\beta$ |
|---|---|---|---|---|---|---|---|
| **Rank** | 1 | 2 | 3 | 4 | 5 | 6 | 7 |
| **K-S statistic** | 0.35 | 0.27 | 0.22 | 0.14 | 0.09 | 0.07 | 0.05 |

#### 2.5 Summary table of incidence of neutropenia for all simulated avadomide dosing schedules.

This section details the full simulation results for all the tested dose and schedule combinations Table S.II. Note that the first 28 days post avadomide administration is the period considered for the observation of neutropenia onset, while ANC profiles were considered up to day 35 for the determination of seven-day neutropenia. This means, for example, that a virtual patient having neutropenia onset at day 27 would be considered towards the toxicity incidence count. Similarly, if that virtual patient maintains toxicity at least up to day 34.5, it would be counted for neutropenia lasting at least seven days. In contrast, a virtual patient experiencing neutropenia onset at day 29 or later would not be considered.

*Table S.II. Full simulation results*

| Schedule | Dose [mg] | Gr3 single [%] | Gr4 single [%] | Gr3 7 days [%] | Gr4 7 days [%] | Recovered Gr3 to above Gr2 [%] | Recovered Gr4 to above Gr2 [%] | Mean time to recover from Gr3 to above Gr2 [day] | Mean time to recover from Gr4 to above Gr2 [day] | AUC [ng/ml*h] | $C_{max}$ [ng/ml] |
|---|---|---|---|---|---|---|---|---|---|---|---|
| 3of7 | 2 | 0.3 | 0 | 0 | 0 | 0 | 0 | | | 286 | 46 |
| 3of7 | 3 | 2.4 | 0 | 0.2 | 0 | 0 | 0 | | | 428 | 69 |
| 3of7 | 4 | 5.3 | 0 | 1 | 0 | 0 | 0 | | | 571 | 91 |
| 3of7 | 5 | 8.3 | 0 | 1.7 | 0 | 0.3 | 0 | 3.28 | | 714 | 114 |
| 3of7 | 6 | 11.5 | 0 | 3.1 | 0 | 1.3 | 0 | 3.46 | | 857 | 137 |
| 3of7 | 7 | 14.9 | 0.3 | 4 | 0 | 2.7 | 0 | 3.3 | | 999 | 160 |
| 3of7 | 8 | 19.2 | 1.5 | 5 | 0 | 5.2 | 0 | 3.39 | | 1142 | 183 |
| 5of7 | 2 | 5.5 | 0 | 2.7 | 0 | 0 | 0 | | | 472 | 48 |
| 5of7 | 3 | 13.5 | 0.2 | 5.4 | 0 | 0 | 0 | | | 709 | 72 |
| 5of7 | 4 | 25.9 | 3.9 | 8.9 | 0 | 0 | 0 | | | 945 | 96 |
| 5of7 | 5 | 36.7 | 6.5 | 13.2 | 0.2 | 1 | 0 | 2.69 | | 1181 | 119 |
| 5of7 | 6 | 45.8 | 9.6 | 20.4 | 1.8 | 0.8 | 0 | 2.74 | | 1417 | 143 |
| 5of7 | 7 | 53.9 | 12.4 | 27.3 | 4.1 | 0.5 | 0 | 2.43 | | 1653 | 167 |
| 5of7 | 8 | 59.7 | 15.7 | 33.7 | 5.4 | 0 | 0 | | | 1889 | 191 |
| 7of14 | 2 | 2.4 | 0 | 0.3 | 0 | 0.2 | 0 | 4.55 | | 336 | 48 |
| 7of14 | 3 | 8.9 | 0 | 1.1 | 0 | 3.8 | 0 | 4.86 | | 504 | 72 |
| 7of14 | 4 | 19 | 2.6 | 3.3 | 0 | 12.5 | 0 | 4.67 | | 672 | 96 |
| 7of14 | 5 | 30.6 | 4.8 | 5.1 | 0 | 23.6 | 0 | 4.66 | | 840 | 120 |
| 7of14 | 6 | 39 | 7.3 | 6.7 | 0 | 31.4 | 0.8 | 4.94 | 6.7 | 1008 | 144 |
| 7of14 | 7 | 46.2 | 9.7 | 8.5 | 0 | 38.4 | 2.3 | 5.16 | 6.34 | 1176 | 168 |
| 7of14 | 8 | 51.9 | 11.9 | 11.1 | 0 | 43.6 | 3.8 | 5.38 | 6.2 | 1344 | 192 |
| 14of28 | 2 | 5.4 | 0 | 1.2 | 0 | 1.4 | 0 | 8.32 | | 338 | 49 |
| 14of28 | 3 | 18 | 2.2 | 4.9 | 0 | 12.6 | 0 | 6.12 | | 507 | 73 |
| 14of28 | 4 | 33.7 | 5.9 | 9 | 0.5 | 28 | 1.4 | 6.26 | 9.51 | 676 | 97 |
| 14of28 | 5 | 45.4 | 10 | 13.1 | 1.1 | 39.4 | 4.6 | 6.54 | 8.15 | 845 | 122 |
| 14of28 | 6 | 55.2 | 14.5 | 18 | 2.3 | 49 | 8.8 | 6.72 | 7.91 | 1014 | 146 |
| 14of28 | 7 | 59.7 | 18.6 | 22.2 | 3.5 | 51.2 | 11.8 | 7.02 | 8.02 | 1183 | 171 |
| 14of28 | 8 | 64.8 | 22.1 | 27.1 | 4.4 | 51.8 | 13.9 | 7.11 | 8.06 | 1352 | 195 |
| 21of28 | 2 | 10.7 | 0 | 6.7 | 0 | 4.5 | 0 | 9.35 | | 505 | 49 |
| 21of28 | 3 | 28.5 | 4.6 | 20.4 | 2.9 | 22 | 0 | 10.2 | | 758 | 73 |
| 21of28 | 4 | 45.4 | 9.2 | 36.6 | 6.8 | 38.5 | 2.4 | 11.24 | 11.71 | 1010 | 98 |
| 21of28 | 5 | 58 | 14.5 | 49.4 | 11.3 | 50.6 | 7.1 | 11.78 | 11.57 | 1263 | 122 |
| 21of28 | 6 | 65.7 | 21.8 | 58.2 | 16.2 | 53.5 | 12.8 | 12.11 | 11.59 | 1515 | 147 |
| 21of28 | 7 | 71.5 | 27.5 | 63.9 | 21.2 | 44.3 | 12.8 | 11.95 | 11.45 | 1768 | 171 |
| 21of28 | 8 | 75 | 31.7 | 68.5 | 25.4 | 28.9 | 8.9 | 11.92 | 11.82 | 2020 | 195 |
| 28of28 | 2 | 11.2 | 0 | 10.9 | 0 | 0 | 0 | | | 652 | 49 |
| 28of28 | 3 | 29 | 4.6 | 28.3 | 4.5 | 0 | 0 | | | 978 | 73 |

| | | | | | | | | | |
|---|---|---|---|---|---|---|---|---|---|
| 28of28 | 4 | 45.9 | 9.6 | 45.6 | 9.1 | 0 | 0 | 1304 | 98 |
| 28of28 | 5 | 58.5 | 14.8 | 57.8 | 14.4 | 0 | 0 | 1630 | 122 |
| 28of28 | 6 | 66.4 | 22.3 | 65.7 | 21.7 | 0 | 0 | 1956 | 147 |
| 28of28 | 7 | 71.9 | 27.7 | 71.4 | 27.4 | 0 | 0 | 2281 | 171 |
| 28of28 | 8 | 75.5 | 32.1 | 75.2 | 31.8 | 0 | 0 | 2607 | 195 |
